# SARS-CoV-2 inhibition in human airway epithelial cells using a mucoadhesive, amphiphilic chitosan that may serve as an anti-viral nasal spray

**DOI:** 10.1101/2020.12.10.413609

**Authors:** Krzysztof Pyrć, Aleksandra Milewska, Emilia Barreto Duran, Paweł Botwina, Rui Lopes, Alejandro Arenas-Pinto, Moutaz Badr, Ryan Mellor, Tammy L. Kalber, Delmiro Fernandes-Reyes, Andreas G. Schätzlein, Ijeoma F. Uchegbu

**Affiliations:** Laboratory of Virology and ABSL3 Animal Facility at the Malopolska Centre of Biotechnology, Jagiellonian University, Gronostajowa 7a, 30-387 Krakow, Poland; Nanomerics Ltd., 6th Floor, 2 London Wall Place, London EC2Y 5AU United Kingdom; UCL School of Pharmacy, 29-39 Brunswick Square, London WC1N 1AX; Centre for Clinical Research in Infection and Sexual Health, UCL Institute for Global Health, Mortimer Market Centre, off Capper Street, London WC1E 6JB; MRC-Clinical Trials Unit at UCL, Institute of Clinical Trials and Methodology, 90 High Holborn, London WC1V 6LJ; Centre for Advanced Biomedical Imaging (CABI), Division of Medicine, University College London, London WC1E 6DD; Computer Science Department, University College London, Gower Street, London WC1E 6BT

**Keywords:** SARS-CoV-2, chitosan amphiphile, N-palmitoyl-N-monomethyl-N, N-dimethyl-N,N, N-trimethyl-6-O-glycolchitosan, GCPQ, anti-viral, viral inhibitor

## Abstract

There are currently no cures for coronavirus infections, making the prevention of infections the only course open at the present time. The COVID-19 pandemic has been difficult to prevent, as the infection is spread by respiratory droplets and thus effective, scalable and safe preventive interventions are urgently needed. We hypothesise that preventing viral entry into mammalian nasal epithelial cells may be one way to limit the spread of COVID-19. Here we show that N-palmitoyl-N-monomethyl-N,N-dimethyl-N,N,N-trimethyl-6-O-glycolchitosan (GCPQ), a positively charged polymer that has been through an extensive Good Laboratory Practice toxicology screen, is able to reduce the infectivity of SARS-COV-2 in A549^ACE2+^ and Vero E6 cells with a log removal value of −3 to −4 at a concentration of 10 – 100 μg/ mL (p < 0.05 compared to untreated controls) and to limit infectivity in human airway epithelial cells at a concentration of 500 μg/ mL (p < 0.05 compared to untreated controls). GCPQ is currently being developed as a pharmaceutical excipient in nasal and ocular formulations. GCPQ’s electrostatic binding to the virus, preventing viral entry into the host cells, is the most likely mechanism of viral inhibition. Radiolabelled GCPQ studies in mice show that at a dose of 10 mg/ kg, GCPQ has a long residence time in mouse nares, with 13.1% of the injected dose identified from SPECT/CT in the nares, 24 hours after nasal dosing. With a no observed adverse effect level of 18 mg/ kg in rats, following a 28-day repeat dose study, clinical testing of this polymer, as a COVID-19 prophylactic is warranted.

## Introduction

There are currently no cures for a wide variety of viral infections, including the ones caused by emerging flaviviruses and coronaviruses, which are regularly causing local outbreaks, epidemics, and pandemics ^1^. Respiratory infections seem to be of special importance, as due to the transmission route, it is almost impossible to control the spread in the population. While common respiratory infections are frequently neglected, it should be borne in mind that seasonal influenza virus claims 200,000 – 500,000 lives annually ^2^. The year 2020 brought us the third zoonotic coronavirus in the 21^st^ century – SARS-CoV-2, causing the COVID-19 disease ^1^. COVID-19 ranges from mild, self-limiting respiratory tract illness to severe progressive viral pneumonia, multiorgan failure and death ^1^. By the beginning of the year the race to develop drugs or re-purpose drugs to prevent coronaviral infection or ease the symptoms had started. Unfortunately, most of the efforts were futile, and the most promising agents such as remdesivir ^3^ and convalescent plasma ^4^ did not live up to their expectations. While passive and active immunisation efforts are ongoing, there is still a need for novel prophylaxis interventions, especially as some of the more promising vaccine technologies neutralise systemic virus, but not viral particles within the nasal epithelia, making it impossible to tell if vaccinated persons would not still transmit the disease ^5^. Here, we describe the activity of a polymer that has been developed previously as a pharmaceutical excipient, has been through a Good Laboratory Practice (GLP) toxicology screen ^6–9^ and may readily be used to prevent or limit COVID-19 infections, e.g., in health-care workers or other persons at risk from severe disease.

Viral binding to cell-surface receptors present in the respiratory tract is a critical step, that enables SARS-CoV-2 to enter the cell and initiate replication. SARS-CoV-2 utilizes the receptor binding domain (RBD) on the spike (S) protein to bind to the angiotensin converting enzyme 2 (ACE2) receptor ^10–12^ present, amongst others, on the ciliated cells of the human respiratory epithelium ^13,14^. This enables activation of the S protein by the cell surface serine proteases and subsequent conformational change, which results in membrane fusion of the viral particle with the cell and ultimately RNA delivery to the replication site ^10,12,15,16^. Blockade of these early steps is a known strategy to prevent the infection of mammalian cells, and has been proven effective for e.g., neutralizing antibodies ^17^ and fusion inhibitors ^18^.

Polymers, such as sulphated glycopolymers have been shown to inhibit the viral binding of human papilloma virus to cell surface receptors ^19^. Sulphated chitosan compounds (i.e. N-carboxymethylchitosan-N,O-sulfate) have been found to inhibit the synthesis of virus-specific proteins and the replication of HIV-1 in cultured T-cells as well as the replication of the Rausher murine leukemia virus in cultured mouse fibroblasts ^20^. Additionally 6-deoxy-6-bromo-N-phthaloyl chitosan ^21^ and chitosan itself ^22–26^ have been reported to have antiviral activity via a variety of mechanisms. It is well known that quaternary ammonium compounds (QACs) are viricidal due to mechanisms involving virion degradation and nucleic acid binding 27. However WO2013/172725 ^28^ reports that N-(2-hydroxypropyl)-3-trimethylammonium chitosan chloride (HTCC, Figure 1), a chitosan QAC with a molecular weight of 50 – 190 kDa (based on viscosity) ^29,30^, and a level of quaternary ammonium groups ranging from 57% to 77% ^31^, inhibits coronavirus infections (e.g. HCoV-NL63) *in vitro* by a mechanism that involves an inhibition of viral entry into the cell ^31^. Positively charged HTCC was shown to electrostatically bind the coronaviral S proteins, blocking its interaction with the entry receptor and consequently the virus replication ^30–32^. Such activity was shown for several members of the *Coronaviridae* family and HTCC also inhibited entry of highly pathogenic SARS-CoV-2 and MERS-CoV into cells ^33^. The HTCC variant effective for the SARS-CoV-2 has a relatively high molecular weight (50 - 190 kDa) ^29,30^, a high level of quaternary ammonium group substitution (57 – 77mole%) ^33^ and has not been through a GLP toxicology screen.

**Figure 1:**
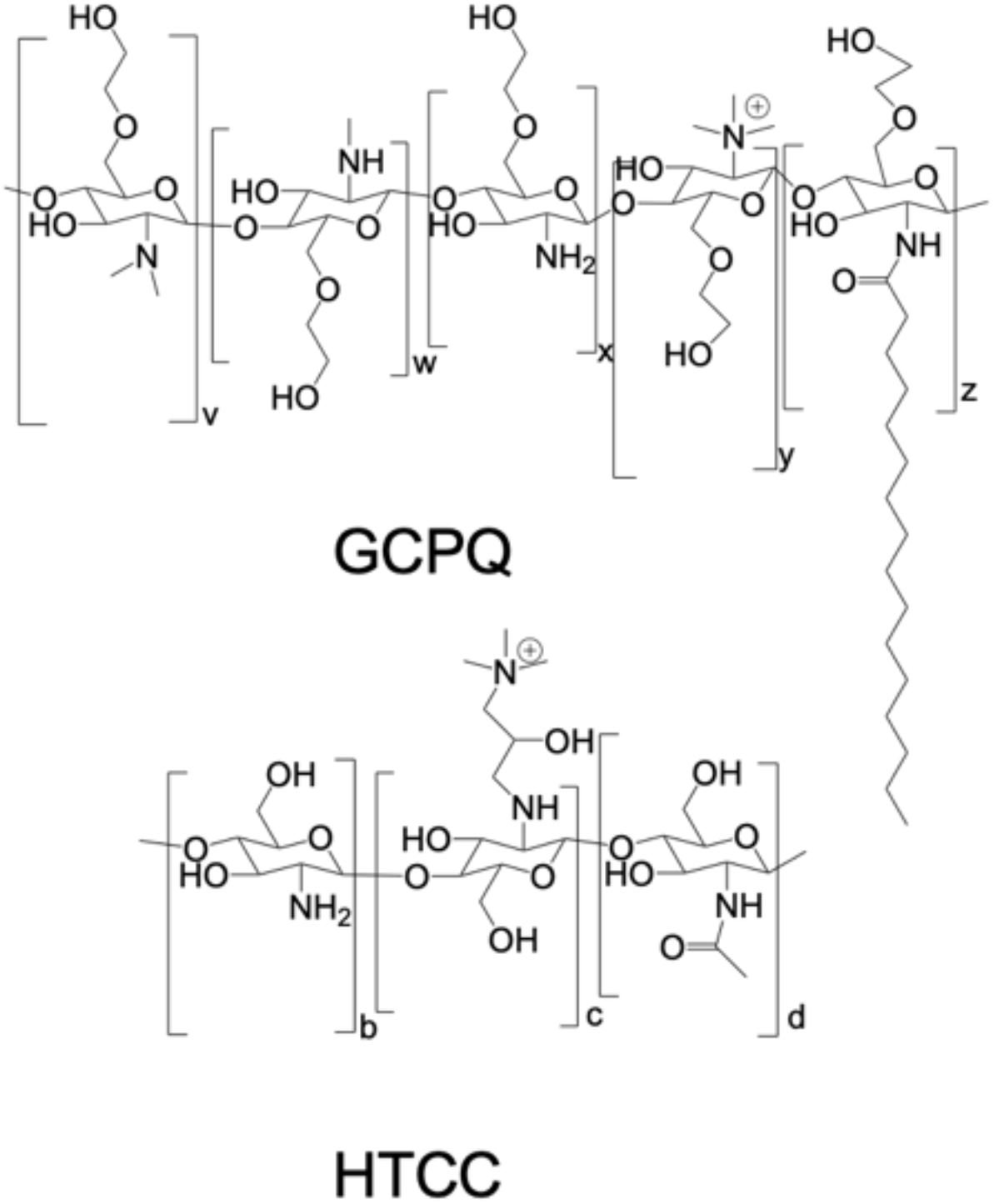
GCPQ and HTCC

Here we show that a different quaternary ammonium chitosan (N-palmitoyl-N-monomethyl-N,N-dimethyl-N,N,N-trimethyl-6-O-glycolchitosan – GCPQ ^6^, Figure 1), with a 6-O-glycol group (lacking in HTCC), a hydrophobic acyl group (lacking in both HTCC and HM-HTCC; the latter derivatised with N-dodecyl groups), a lower molecular weight (10 – 30 kDa) than HTCC, a trimethyl quaternary ammonium group directly in place of the C2 amine group in chitosan, unlike HTCC (which has the hydroxypropyltrimethylammonium group attached to the C2 nitrogen), and a lower level of quaternary ammonium substitution than HTCC (less than 40 mole%) is also able to inhibit viral entry into cells. Oligochitosans without the quaternary ammonium group were inactive in inhibiting coronavirus entry into cells ^30^. However the quaternary ammonium group on HTCC is not the only structural requirement for activity, as many quaternary ammonium polymers were found to be inactive in inhibiting coronavirus entry into cells ^30^. Furthermore HTCC was inactive in inhibiting a number of other viruses (e.g. human herpes virus 1, influenza A, adenoviruses and enteroviruses) ^30^. Molecular weight determinants of activity are also unclear, as while high molecular weight HTCCs (50 - 190 kDa) ^29,30^ were active against coronaviruses, chitosans of molecular weight 5 – 17 kDa were more effective antiviral agents against tobacco mosaic virus in Xanthi-nk tobacco leaves (viral inhibition of 58 – 87%) than chitosans with a molecular weight of 130 kDa and above ^25^. It is thus clear that it is not straightforward to define the polymer structure features that will inhibit coronaviruses in cells, or indeed inhibit a broad spectrum of viruses.

We decided to study GCPQ’s anti-viral properties, as crucially, GCPQ is being developed as a pharmaceutical excipient and has been through a Good Laboratory Practice (GLP) toxicology screen ^6–9,34,35^, with Investigational New Drug enabling studies currently ongoing and funded by the US National Institute of Health National Center for Advancing Translational Sciences (NCATS). The existing GCPQ safety data and the anti-viral activity, reported here, strongly favour the clinical testing of GCPQ as an anti-viral nasal spray.

## Materials and Methods

### Materials

Vero E6 (Cercopithecus aethiops; kidney epithelial; ATCC: CRL-1586) and A549 cells with ACE2 overexpression (A549^ACE2+^) ^33^ were used in the study. For all cultures Dulbecco’s MEM (ThermoFisher Scientific, Poland) supplemented with 3% foetal bovine serum (heat-inactivated; ThermoFisher Scientific, Poland) and antibiotics: penicillin (100 U/ml), streptomycin (100 μg/ mL), and ciprofloxacin (5 μg/ml) were used. Commercially available MucilAir HAE cultures were used for the *ex vivo* analysis (Epithelix Sarl, Switzerland). All cultures were carried out at 37°C under 5% CO_2_.

SARS-CoV2 was isolate 026V-03883 (Charité – Universitätsmedizin Berlin, Germany, European Virus Archive - Global - EVAG, https://www.european-virus-archive.com).

The XTT cell viability kit (Biological Industries, Israel) was used for the cell viability assays. The following reagents were also used: viral DNA/RNA isolation kit (A&A Biotechnology, Poland), High-capacity cDNA reverse transcription kit (Thermo Fisher Scientific, Poland), Real-time qPCR kit (RT-HS-PCR mix probe, A&A Biotechnology, Poland) and the real-time qPCR oligonucleotides are listed in Table 1.

**Table 1:**
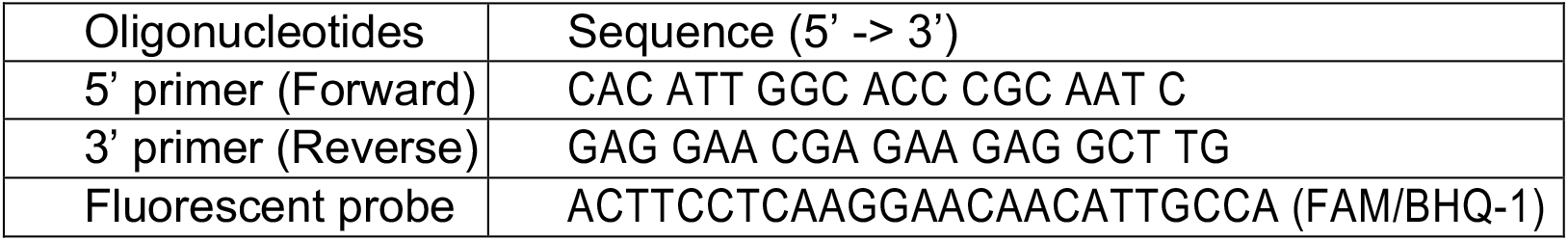
Real-time qPCR oligonucleotides.

The GPCQ compounds are listed in Table 2. The compounds were suspended in 1 × PBS to the final concentration of 5 mg/ mL. All stocks were stored at 4°C until use.

**Table 2:** GCPQ samples

|  |  |
| --- | --- |
| GCPQa | GCPQc |
| Molecular Weight = 10 kDa | Molecular Weight = 15 kDa |
| Mole% palmitoyl groups = 18 | Mole% palmitoyl groups = 18 |
| Mole% quaternary ammonium groups = 18 | Mole% quaternary ammonium groups = 20 |
| GCPQb | GCPQd |
| Molecular Weight = 30 kDa | Molecular Weight = 60 kDa |
| Mole% palmitoyl groups = 19 | Mole% palmitoyl groups = 18 |
| Mole% quaternary ammonium groups = 19 | Mole% quaternary ammonium groups = 16 |

### Methods

#### Tissue culture

The cytotoxicity of compounds was assessed by incubating confluent monolayers of Vero E6 and A549^ACE2+^ cells with a range of GCPQ compound concentrations. The XTT assay was carried out 48 hours later, according to the manufacturer’s protocol using 200,000 cells per well and DMEM supplemented with foetal bovine serum, penicillin and streptomycin (please see above under materials).

The ability of each compound to inhibit the virus replication was determined by infecting confluent Vero E6 and A549^ACE2+^ monolayers with the SARS-CoV-2 virus at 400 TCID_50_/mL in the presence of test compounds or phosphate buffered saline - PBS (TCID_50_ = 50% Tissue Culture Infectious Dose). Mock controls (cell lysate without the virus) and medium (supplemented DMEM – please see above) controls were included. Each compound was present during and after the infection. The cells were then incubated for 2 hours at 37°C and 5% CO_2_. Afterward, the cells were washed three times with PBS, and each compound was re-applied onto the cell monolayer. 100μL of the cell culture supernatants were subsequently collected from each designated well after two days of culture. The experiments were carried out in triplicate.

Virus replication inhibition in HAE was evaluated by infecting MucilAir™ (Epithelics Sarl, Switzerland) with SARS-CoV-2 virus at 5000 TCID_50_/mL in the presence of GCPQa or PBS. GCPQa diluted in PBS was added to the apical side of the insert (200 μg/ml or 500 μg/ml) and incubated at 37°C for 30 minutes before the infection. After the pre-incubation was completed, the compound was removed and fresh dilutions of the compound with the virus were added and incubated for 2 hours at 37°C. Next, the apical side of the HAE was washed thrice with PBS and each compound was re-applied and incubated again for 30 minutes at 37°C. After the last incubation with the GCPQ, the samples (50 μL) were collected and the HAE cultures were left in air-liquid interphase. Every 24 hours the HAE apical surface was incubated for 30 minutes with the GCPQ or PBS and the samples were collected for virus yield evaluation. Viral RNA was isolated from the apical washings or cell culture supernatant, RNA was isolated (Viral DNA/RNA; A&A Biotechnology, Poland), reverse-transcribed into cDNA (High Capacity cDNA Reverse Transcription Kit; ThermoScientific, Poland), and subjected to the qPCR analysis. Briefly, cDNA was amplified in a reaction mixture containing 1 × qPCR Master Mix (A&A Biotechnology, Poland), in the presence of probe (100nM) and primers (450nM each), sequences provided in Table 1.

The reaction was carried out using the 7500 Fast Real-Time PCR machine (Life Technologies, Poland) according to the scheme: 2 min at 50°C and 10 min at 92°C, followed by 40 cycles of 15 s at 92°C and 1 min at 60°C. In order to assess the copy number for N gene, DNA standards were prepared, as described before ^36^. The obtained data is presented as virus yield and as the log removal value (LRV), showing the relative decrease in the amount of virus in cell culture media compared to the control.

### Intranasal Delivery in a Healthy Animal Model

GCPQ (molecular weight = 10 kDa, mole% palmitoyl groups = 16 and mole% quaternary ammonium groups = 13) was radiolabelled using a two stage strategy: first an acylating reagent [N-succinimidyl-3[4-hydroxyphenyl]propionate - the Bolton and Hunter reagent (BH)] was initially covalently coupled to GCPQ and then the GCPQ-BH complex was iodinated with ^125^I. Briefly, GCPQ (90 mg) was dissolved in DMSO (3 mL). To this solution was added 200 μL of triethylamine and 0.05 molar equivalents (10 mg) of BH reagent and the reaction allowed to proceed overnight at room temperature with stirring. The next day, the GCPQ-BH conjugate was precipitated using an acetone: diethyl ether mixture (1:2, v/v) and the pellet was wash 3 times with the same acetone: diethyl ether mixture. The washed pellet was dissolved in methanol (2 mL) and dialyzed against water overnight. The dialysed GCPQ-BH was then freeze dried and collected. Labelling of GCPQ-BH with ^125^I was performed using iodination beads ® (Thermo Scientific Pierce, UK). Briefly, GCPQ-BH (20 mg) and 100 mg GCPQ were dissolved in methanol with stirring then the methanol was removed under vacuum and Tris-HCL buffer (25 mM, pH 4.8, 1.8 mL) was added to the dry film to produce a final concentration of 66.7 mg/mL. This solution was then added to a tube containing the I^125^ (1 mCi, 17 Ci/mg, 0.392 nmol, Perkin-Elmer, USA) and four iodination beads® (Thermo Scientific Pierce, UK). The reaction was incubated for 1.5 hours at room temperature, after which the reaction was terminated by separating the solution from beads. PD Spin Trap G-25 Columns (GE Healthcare Life Sciences, UK), that are prepared by vortexing and discarding of the eluting storage buffer by centrifuging (2800 rpm for 1 min), were used in order to remove the free iodine (with the free iodine removed through the addition of 50 μL of the reaction per column and centrifuging at 2,800 rpm for 2 min). The eluent was placed in Amicon ultra centrifugal filters (3 kDa, Millipore, USA) with 200 μL H_2_O, and was subject to repeated washes (through centrifuging at 10,000 rpm for 10 min), until the washed out water produced negligible counts. All animal experiments were performed under a UK Home Office licence and were approved by the local ethics committee. A Male Balb/C mouse weighing 25 g (Charles River, UK), allowed free access to standard rodent chow and water, was intranasally administered radiolabelled GCPQ-BH (10 mg/kg, 1.2 MBq) by using a pipette to place 5uL of the radiolabelled material into the mouse nares and allowing the mouse to sniff in the dose. At various time points after the administration of the radiolabelled GCPQ-BH, animals were anaesthetised using isofluorane (1-2% in oxygen), maintained at 37 °C and submitted for NanoSPECT/CT analysis (Mediso, USA).

### In vivo SPECT/CT Imaging and Analysis

SPECT/CT scans of the mouse head at 30 min, 2h 30 min and 24 h after nasal administration were acquired using a NanoSPECT/CT scanner (Mediso, Hungary). The mouse was <u>anaesthetised using isofluorane (1-2% in oxygen)</u> and maintained at 37°C. SPECT images were obtained over 30 minutes using a 4-head scanner with nine 1.4 mm pinhole apertures in helical scan mode with a time per view of 60 seconds. CT images were subsequently acquired using a 45 kilo volt peak (kVp) X-ray source, 500 ms exposure time in 180 projections, a pitch of 0.5 with an acquisition time of 4:30 minutes. Body temperature was maintained by a warm air blower and the respiration and core body temperature was monitored throughout. CT images were reconstructed using Bioscan InVivoScope (Bioscan, USA) software in voxel size 124 × 124 × 124 μm, whereas SPECT images were reconstructed using HiSPECT (ScivisGmbH, Bioscan) in a 256 × 256 matrix. Images were fused and analysed using VivoQuant (Invicro, A Konica Minolta Company) software. 3D Regions of Interest (ROIs) were created for the uptake within the nares for each time point and the activity calculated as the percentage of administered dose. Representative images are scaled the same (same min and max). After the final scan the mouse was sacrificed and the entire head of the mouse analysed using a curimeter (Capintech, Mirion Technologies, UK) for *ex vivo* validation of ^125^I concentration.

## Results

### Cytotoxicity

Four GCPQ polymers (see Table 2) were tested. First, the cytotoxicity of polymers was analysed on Vero E6 and A549^ACE2+^ cells. The results of the analysis are shown in Figure 2.

**Figure 2:**
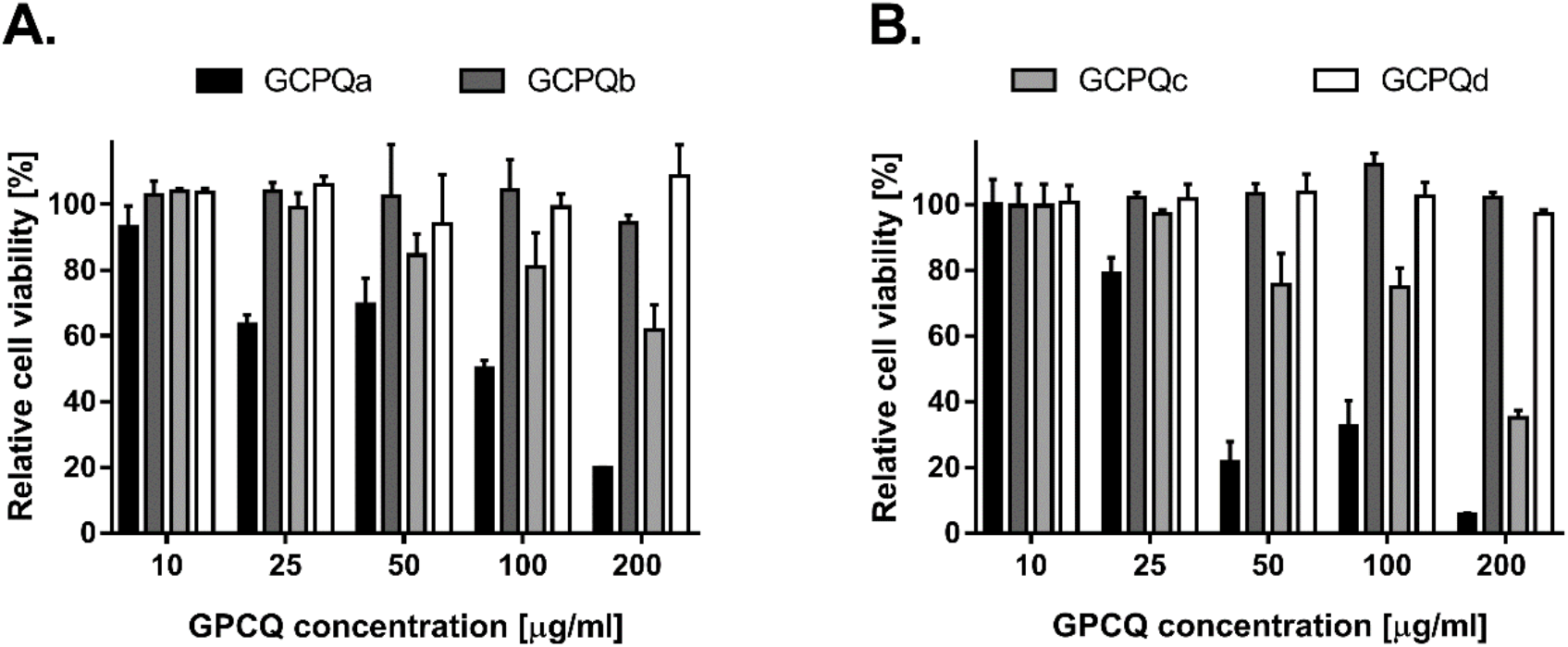
Cytotoxicity of GCPQs *in vitro*. Cell viability was assessed using an XTT assay on Vero E6 cells (A) and A549^ACE2+^ cells (B). Relative viability of cells (percentage of the untreated control) is shown on y-axis. All assays were performed in triplicate, and average values with standard errors are presented. The letters a to d refer to the GCPQs shown in **Table 2**.

For the virus assays, only non-toxic concentrations were tested: 10 μg/ml of GCPQa, 25 μg/ml for GCPQc, and 200 μg/ml for GCPQb and GCPQd.

### Anti-viral activity in Vero E6 and A549 cells

The anti-viral activity of GPCQs was analysed on Vero E6 and A549^ACE2+^ cells. Each analysis was performed in triplicate, and the experiment was repeated twice. The results are presented in Figure 3. The assay showed inhibition of SARS-CoV2 replication in the presence of GCPQa and GCPQc at non-toxic concentrations.

**Figure 3:**
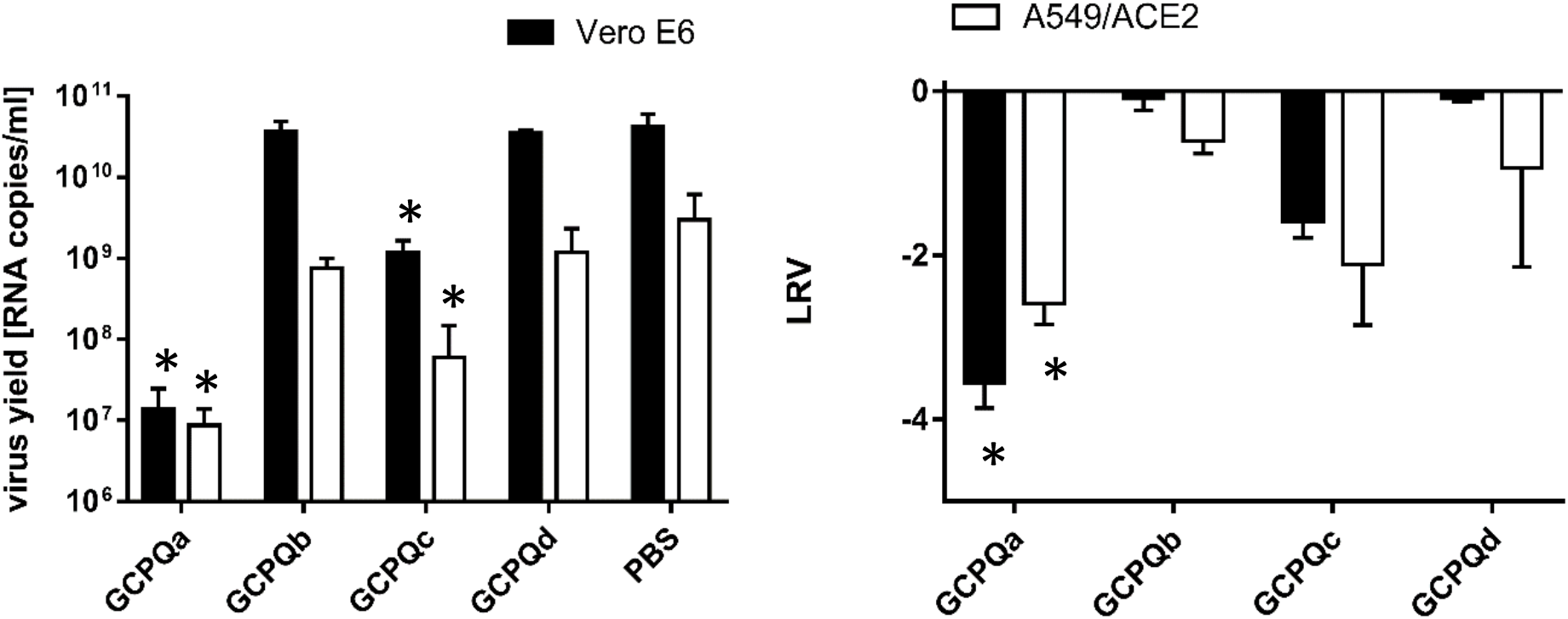
Anti-viral activity of GCPQs against SARS-CoV-2. Virus replication was evaluated using RT-qPCR. The data are presented as a number of RNA copies per mL of the original sample (left) or as Log Removal Value (LRV) compared to untreated samples (right). Non-toxic concentrations were tested (10 μg/ml of GCPQa, 25 μg/ml for GCPQc, and 200 μg/ml for GCPQb and GCPQd). The assay was performed in triplicate, and average values with standard errors are presented. For the RNA copies per mL data GCPQa and GCPQc are significantly different from PBS treated cells (p < 0.0001) and for the log removal value, GCPQa is significantly different from PBS treated samples in the A549 and Vero E6 cell lines (p < 0.05).

Our analysis demonstrated that GCPQa and GCPQc effectively inhibit SARS-CoV-2 replication *in vitro* at non-toxic concentrations. GCPQa showed the highest toxicity, but at the same time highest anti-SARS-CoV-2 potential (~4 logs decrease in viral load at 10 μg/mL).

### Viral inhibition in human airway epithelial (HAE) cells

The effectiveness of GCPQ in the artificial cell culture systems has its drawbacks and for that reason it is of importance to validate the observations in more complex systems that replicate the host-pathogen interactions. For example, we have observed lack of toxicity *in vivo* at much higher concentrations ^7^, while the immortalized cell lines were susceptible even at low micromolar concentrations. This most likely results from the effect of the charged polymers on the cell adhesion to the plastic, which is not relevant in the tissue. Furthermore, chitosan and its derivatives were shown to carry strong antineoplastic properties, and these properties may affect the cell viability in culture.

To better validate our observation on the anti-SARS-CoV-2 activity of GPCQa, a fully differentiated HAE *ex vivo* model was used, reconstituting *ex vivo* the human respiratory epithelium. Two different concentrations of GPCQa were evaluated (200 μg/mL and 500 μg/mL) and PBS was used as a control. Each analysis was performed in triplicate and the results are shown in Figure 4.

**Figure 4:**
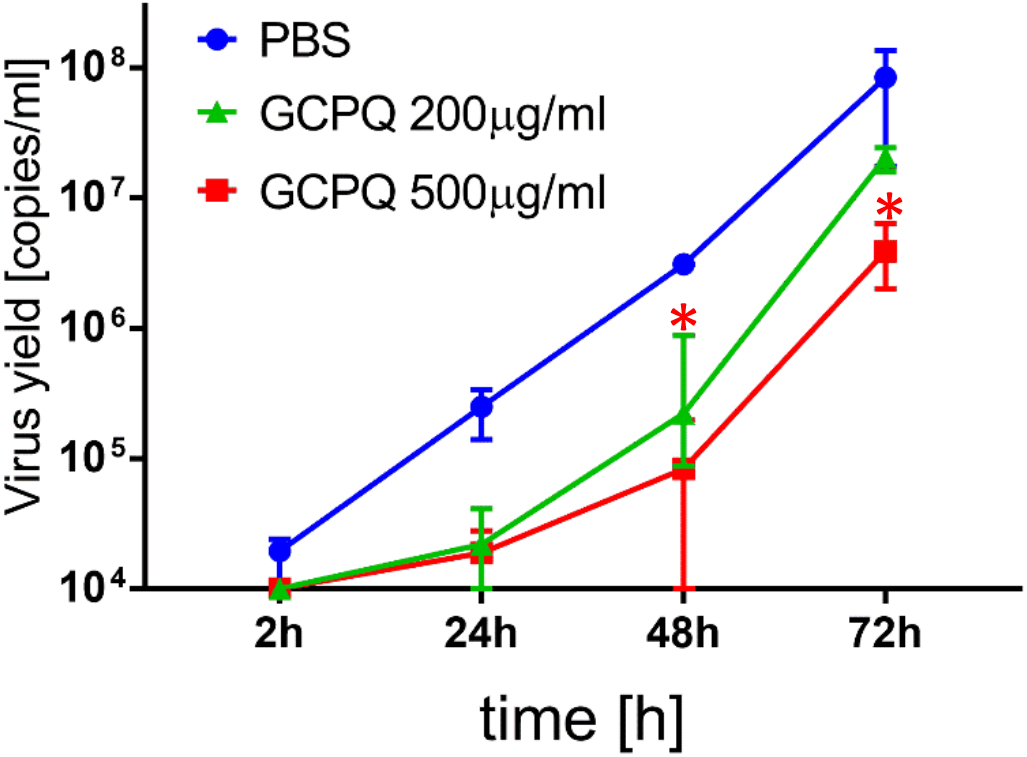
Replication of SARS-CoV-2 in fully differentiated tissue cultures of the human respiratory epithelium (HAE) in the presence or absence of GPCQ. Virus replication was evaluated using RT-qPCR. The data are presented as a number of viral copies per ml. The assay was performed in triplicate, and median values with range are presented. At the 48 and 72 hour time points, GCPQa at 500 μg/ mL is significantly different from PBS treated samples (p < 0.05).

The results show that GCPQa inhibits SARS-CoV-2 replication in the HAE *ex vivo* model. A lower viral yield was detected in the cultures treated with GCPQa than in the control cultures treated with PBS after 72 h of infection.

### Nasal Delivery

The intranasal delivery of GCPQ to mouse nares resulted in the polymer being detectable within the nares area of the mouse head up to 24 hours after dosing as would be expected since the polymer is mucoadhesive ^9^. 3D ROI’s of the GCPQ uptake on SPECT images indicated that directly after administration (30 min) 28.22% of the administered dose was found to be within the nares, after 2h 30 min this had reduced slightly to 25.13% of the administered dose and at 24 h, 13.13% of the administered dose was found to be retained in the nares. *Ex vivo* curimeter analysis of the mouse head at 24 h, also confirmed that 13.5% of the administered dose was still present in the mouse head, which is slightly higher than that obtained by SPECT, but is reflective of the radioactivity within the whole head and not just the nares area.

## Discussion

Here we introduce GCPQ, a low molecular weight chitosan derivative with features that unexpectedly confer anti-viral activity. GPCQ is being developed as a pharmaceutical excipient, has been through a GLP toxicology screen and a no observed adverse effect level (NOAEL) determined for a 28 day repeat dose in the rat (18 mg/ kg) ^7^. It is the key excipient in the enkephalin pain therapeutic being developed by NCATS. Human nasal concentrations in excess of 1mg/ mL will be obtained by dosing 1mg in each nare, a dose of 0.03 mg/ kg that is 600 fold lower than the NOAEL dose. A low molecular weight clearly promotes activity against SARS-COV-2 in mammalian cells (Table 2 and Figures 3 – 4) and this is correlated with the ease with which this polymer may be incorporated into aqueous media. Glycol chitosans of molecular weights 40 kDa and 100 kDa were not active (data not shown), demonstrating that quaternary ammonium and possibly palmitoyl groups are important determinants of activity. We speculate that GCPQ acts by binding to the virus via electrostatic interactions in a similar manner to HTCC ^30–32^.

GCPQ possesses some advantages for use in viral inhibition and specifically the clinical prevention of viral infections as GCPQ is mucoadhesive ^9^, has a long residence time in the nares (Figure 5) and is chemically stable for at least 18 months ^6^. GCPQ also self assembles into nanoparticles and these nanoparticles may be clustered into microparticles for nasal delivery ^7^, as required by the regulator ^37^. We hypothesise that GCPQ could be used as a molecular mask nasal spray for the prevention of coronavirus infections. The new data showing that SARS-COV-2’s neurological symptoms (such as loss of smell and taste, headache, fatigue, nausea and vomiting in more than one-third of individuals and impaired consciousness) is correlated with the entry of SARS-COV-2 into the brain via the olfactory neurons, due to presence of the virus in the nasal cavity ^38^, means that local interventions, such as with GCPQ that limit viral cell entry in the nasal cavity could have a profound impact on the course and severity of the disease.

**Figure 5:**
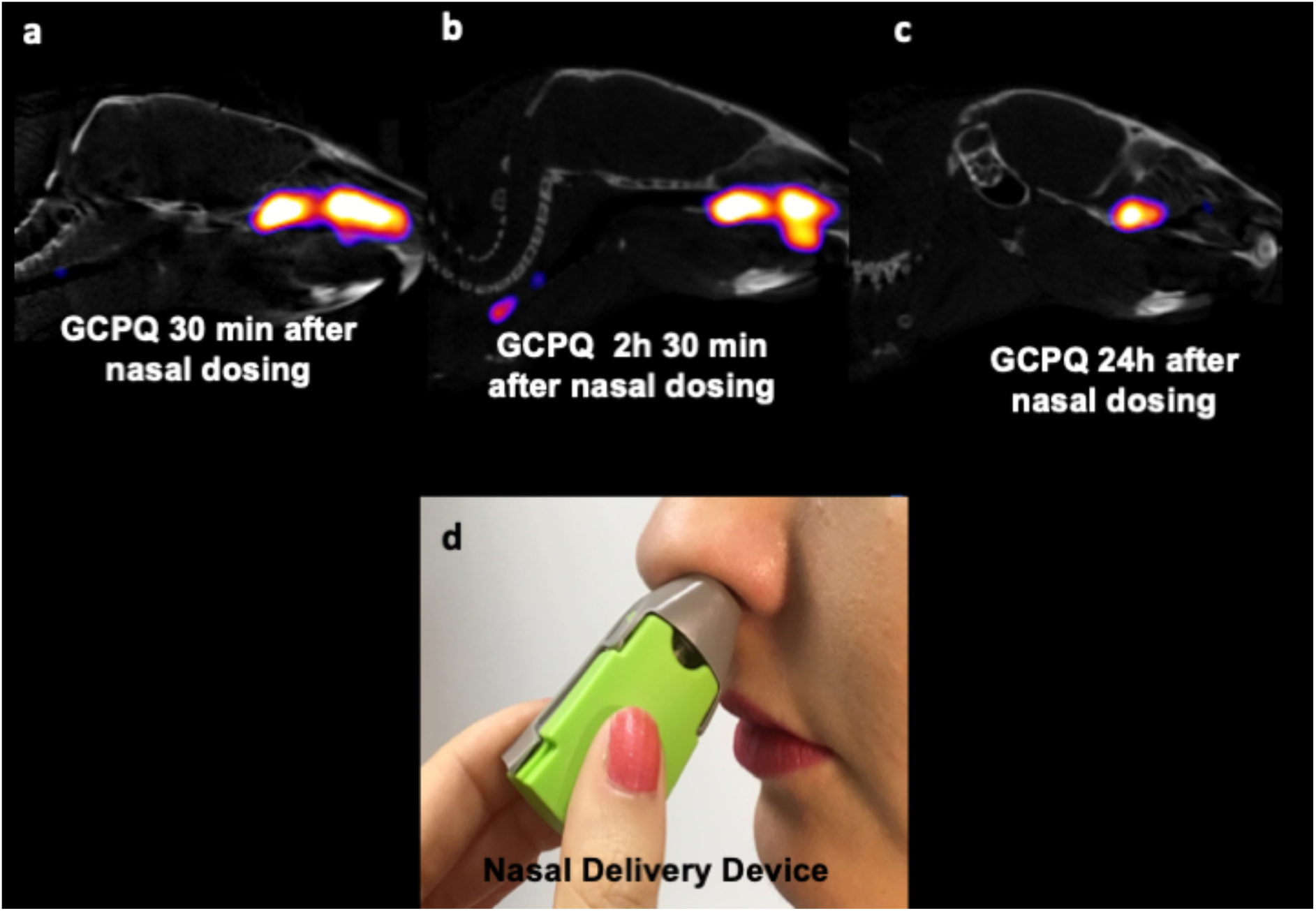
Sagittal SPECT/CT images of radiolabelled GCPQ (10 mg/ kg) at 30 min, 2h 30 min and 24 hours after nasal administration to male mice (a-c), the nasal delivery device that may be used to deliver the prophylactic GCPQ powder (d).

The potential applicability of GCPQ in the prevention of viral infections is supported by the fact that carrageenans (anionic sulphated carbohydrates) have been shown to reduce the duration of disease (reduced by 3 days) in influenza and common cold patients (reduced the number of relapses over a 21 day period by three times) and prevent influenza A viral infections in mice, acting by preventing viral interaction with relevant cell surface receptors ^39,40^.

As GCPQ’s activity is not predicated specifically on the recognition of particular epitopes but appears to be based on electrostatic interactions between GCPQ and the virus, it is possible that GCPQ may be applied to a wide variety of viral infections. These advantages mean that GCPQ may be given as a nasal spray or by other means for the prevention and treatment of specific viral infections.

## Conclusions

A mucoadhesive polymer with anti-SARS-COV-2 activity is presented. The polymer may be used as a nasal spray to prevent SARS-COV-2 infections.

## Acknowledgements

None

## Author Contributions

IFU and AGS conceived of the study and IFU drafted the manuscript. KP, AM, EBD and PB carried out the *in vitro* tests and KP contributed to the writing of the manuscript. RL and TLK carried out the animal studies. MB synthesised the GCPQ, RM synthesised glycol chitosan and provided information on GCPQ, AAP and DFR contributed to the discussion in the manuscript.

